# Measurement invariance explains the universal law of generalization for psychological perception

**DOI:** 10.1101/341305

**Authors:** Steven A. Frank

## Abstract

The universal law of generalization describes how animals discriminate between alternative sensory stimuli. On an appropriate perceptual scale, the probability that an organism perceives two stimuli as similar typically declines exponentially with the difference on the perceptual scale. Exceptions often follow a Gaussian probability pattern rather than an exponential pattern. Previous explanations have been based on underlying theoretical frameworks such as information theory, Kolmogorov complexity, or empirical multidimensional scaling. This article shows that the few inevitable invariances that must apply to any reasonable perceptual scale provide a sufficient explanation for the universal exponential law of generalization. In particular, reasonable measurement scales of perception must be invariant to shift by a constant value, which by itself leads to the exponential form. Similarly, reasonable measurement scales of perception must be invariant to multiplication, or stretch, by a constant value, which leads to the conservation of the slope of discrimination with perceptual difference. In some cases, an additional assumption about exchangeability or rotation of underlying perceptual dimensions leads to a Gaussian pattern of discrimination, which can be understood as a special case of the more general exponential form. The three measurement invariances of shift, stretch, and rotation provide a sufficient explanation for the universally observed patterns of perceptual generalization. All of the additional assumptions and language associated with information, complexity, and empirical scaling are superfluous with regard to the broad patterns of perception.

## Introduction

The probability that an organism perceives two stimuli as similar typically decays exponentially with separation between the stimuli. The exponential decay in perceptual similarity is often referred to as the universal law of generalization (Shepard, 1987; Chater & Vitányi, 2003).

*Generalization* arises because perceived similarity may describe recognition of a general category. For example, two circles may have different sizes, colors, and shadings. Perceived similarity arises from the generalized perception of *circle* as a category.

*Universal law* arises because many empirical observations fít the pattern for diverse sensory modalities across different species. Typical exceptions take on a Gaussian probability pattern for perceived separation (Ghirlanda & Enquist, 2003).

Both theory and empirical analysis depend on the definition of the perceptual scale. How does one translate the perceived differences between two circles with different properties into a quantitative measurement scale?

There are many different suggestions in the literature for how to define a perceptual scale. Each of those suggestions develop very specific notions of measurement based, for example, on information theory, Kolmogorov complexity theory, or multidimensional scaling descriptions derived from observations (Shepard, 1987; Chater & Vitányi, 2003; Sims, 2018).

I focus on the minimal properties that any reasonable perceptual measurement scale must have rather than on detailed assumptions motivated by external theories of information, complexity, or empirical scaling. I express the minimal properties as simple invari ances.

I show that a few inevitable invariances of any reasonable perceptual scale determine the exponential form for the universal law of generalization in perception. All of the other details of information, complexity, and empirical scaling are superfluous with respect to understanding why the universal law of generalization has the exponential form.

I also show that, when the separation between stimuli depends on various underlying perceptional dimensions, it sometimes makes sense to assume that the perceptual scale will also obey exchangeability or rotational invariance. When that additional invariance holds, the universal law takes on the Gaussian form, which I will show to be a special case of the general exponential form.

## Basic problem and notation

Chater and Vitányi Chater & Vitányi (2003) state the law as “the probability of perceiving similarity or analogy between two items, *a* and *b*, is a negative exponential function of the distance *d*(*a, b*) between them in an internal psychological space.”

Let the notation *P*(*R_b_* |*S_a_*) describe the probability of a positive response, *R_b_*, to the event *b*, given an initial stimulus, *S_a_*, by the event *a*. A positive response expresses the perceived similarity of *b* to *a*, which may also be thought of as expressing the generalization that *b* and *a* belong to the same category.

The goal here is to understand how the perceived similarity of *b* to *a*, observed as *R_b_* |*S_a_*, translates into a continuous psychological measurement scale, *T_b|a_*, so that

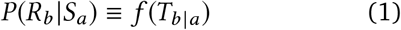

for a suitably defined mapping *R_b_*|*S_a_* ↦ *T_b |a_* and probability distribution function, *f*. We seek the characteristics of the mapping and the associated function, *f*.

## Invariant properties of measurement

There are many different suggestions in the literature for how to define a perceptual scale, *T_b|a_* (Shepard, 1987; Chater & Vitányi, 2003; Sims, 2018). I focus on the minimal properties that any reasonable measurement scale must have, rather than on detailed assumptions motivated by external theories (Luce & Narens, 2008; Narens & Luce, 2008; Houle et al., 2011). I express the minimal properties as simple invariances. Before listing the invariances, consider two simple examples.

First, suppose we wish to analyze the perception of temperature for event *b*, given that event *a* is at the freezing point for water. If we choose to measure the temperature on the Celsius scale, then *T_a|a_* = 0 and *T_b|a_* = *C*. It would make sense to assume that perceptual generalization would be identical if we assigned numerical values on a Fahrenheit scale, 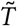, which we obtain by 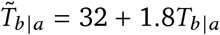.

Second, suppose we wish to measure the perception of separation between two potentially dangerous prey items, such as noxious butterflies Sims (2018); Brower (1958). We begin by exposing a noxious butterfly, *a*, to a predator. After the predator tastes butterfly *a*, we then expose butterfly *b* to the same predator. For the exposure to *b*, we measure the tendency for the predator to attack the potential prey item. Data may include the directions of movements relative to the butterfly, attacks per minute, or the probability of attack over repeated experiments. We now wish to find *a* scale, *T_b|a_*, that is a function of the data we have for the response to various butterflies, *b*, relative to an initial stimulus butterfly, *a*.

However we choose that scale, it makes sense to suppose that the information in *T_b |a_* about the perceptual separation between b and a is the same as the information in *α* + *βT_b|a_* for some constants *α* and *β*. If that were not so, it would be equivalent to saying that the analogs of Celsius and Fahrenheit scalings would provide different information about the perceptual separation between the two butterflies.

For example, we may wish to set *T_a|a_* = 0 to describe a zero separation between identical butterflies, or we may wish to let *T_a|a_* express the amount of the baseline predator perception of the separation between identical stimuli. In either case, our scale *T* should contain the same information with respect to the probabilities of response given in eqn 1. Here, similarity associates with the probability of avoidance response. We may also wish to express our scale standardized with respect to a unit response, *T_b^*^|a_*, to *b*\*, or with respect to a unit response to *T_b^†^|a_*, to *b^†^*. The constant multiplications required to transform between units of measure should not alter the information in the perceptual scale, T, about the probabilities of response.

## Affine and rotational invariance

In other words, the way in which we measure perceptual distance between two stimuli should be independent of a shift and stretch of the scale by constant values. Formally, the scale should be shift invariant with respect to any constant, *α*, such that

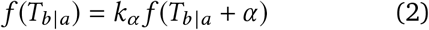

for some constant of proportionality, *k_α_*. The scale should also be stretch invariant to any constant, *β*, such that

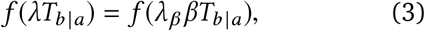

for which I show below that *λ = λ_β_β* is an invariant constant that is conserved in any particular application, set by the fact that 1/*λ* is the average value on the perceptual scale for positive responses to varying events *b* for a given stimulus *a*.

Thus, the scale *T_b|a_* has the property that the associated probability pattern is invariant to the affine transformation of shift and stretch, *T_b|a_* ↦ *α* + *βT_b|a_*. I will show that affine invariance by itself determines the exponential form for the universal law of generalization in perception.

In some cases, it makes sense to assume that the perceptual scale should also obey rotational invariance, such that the Pythagorean partition

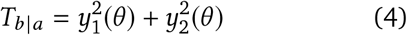

splits the measurement into components that add invariantly to *T_b|a_* for any value of the parameter *θ*. The invariant quantity *T_b|a_* defines a circle in the (*y*_1_, *y*_2_) plane with a conserved radius 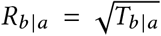 that is invariant to *θ*, the angle of rotation around the circle, circumscribing a conserved area 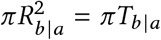.

Rotational invariance partitions a conserved quantity into additive components, for which the order may be exchanged without altering the invariant quantity. When rotational invariance holds, the universal law takes on a Gaussian form, which we will see to be a special case of the general exponential form.

The following sections develop the three invariances of shift, stretch, and rotation. I show that essentially all of the common properties of perceptual generalization follow from these invariances. The analysis here briefly summarizes the detailed development in Frank (2016). The novelty in this article concerns the simple understanding of widely observed psychological patterns.

## Shift invariance implies the exponential form

To simplify notation, denote the perceptual scale by *x* ≡ *T_b|a_* and the associated probability distribution by *f*(*x*) ≡ *f* (*T_b|a_*). If we assume that the functional form for the probability distribution, *f*, is invariant to a constant shift of the perceptual scale, *x* + *α*, then by the conservation of total probability

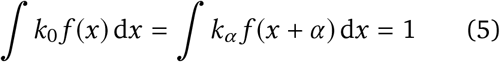

holds for any magnitude of the shift, *α*, in which the proportionality constant, *k_α_*, changes with the magnitude of the shift, *α*, independently of the value of *x*, in order to satisfy the conservation of total probability.

From this equality for total probability, which holds for any shift *α* by adjustment of the constant, *k_α_*, the condition for *x* ≡ *T_b|a_* to be a shift-invariant scale is equivalent to

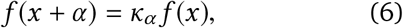

in which *κ_α_* depends only on *α* and is independent of *x*. Because the invariance holds for any shift, *α*, it must hold for an infinitesimal shift, *α* = *ε*. We can write the Taylor series expansion for an infinitesimal shift as

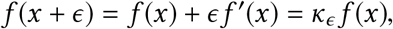

with *κ_ϵ_* = 1 – *λ_ϵ_*, because *ϵ* is small and independent of *x*, and *κ*_0_ = 1. Thus,

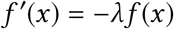

is a differential equation with solution

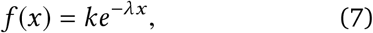

in which *k* is determined by the conservation of total probability. When the perceptual scale ranges over positive values, *x* > 0, then *k* = *λ*.

The assumption that a perceptual scale must be shift invariant is, by itself, sufficient to explain the exponential form of the universal law of generalization.

## The exponential form implies shift invariance

The previous section showed that if the perceptual scale, *x*, is shift invariant, then the exponential form of the universal law of generalization follows. This section shows that if the universal law of generalization takes on the exponential form, then the underlying perceptual scale must be shift invariant. Thus, shift invariance is necessary and sufficient for the exponential form. Any assumptions about the perceptual scale beyond shift invariance must be superfluous with respect to the exponential form.

Begin with the assumption of the exponential form in eqn 7 and write the consequence of a shift of the scale *x* by *a* as

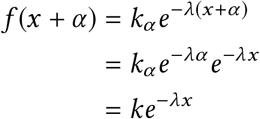

in which *k_α_ = ke^λα^* because the constant multiplier of the exponential must be chosen to satisfy the conservation of total probability, in other words, to normalize the total probability to be one. Thus, the exponential form implies shift invariance of the perceptual scale, *x*.

## Stretch invariance and rate of perceptual change

If we assume that the perceptual scale is defined for positive values, *x* > 0, then the average value of *λx* is always one, because

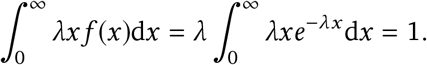

Thus, for average value, 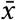, the value of λ is 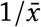. We can think of 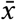 as the average discrimination of various events, *b*, relative to an initial stimulus, *a*, in which the set of events b corresponds to a uniform continuum along the perceptual scale, *x*.

It makes sense to assume that the average discrimination would not change if we arbitrarily multiplied our numerical scale for perception, *x*, by a constant, *β*. The conservation of average value and stretch invariance are equivalent, because

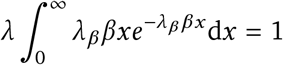

when we allow *λ_β_* to adjust to satisfy the conservation of average value so that *λ* = *λ_β_β* or, equivalently, we assume stretch invariance of the scale, *x* ≡ *T_b|a_*.

The constant 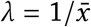 can be thought of as the slope or rate of change in the logarithm of discrimination, because

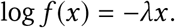

Stretch invariance, or the conservation of average value, is sufficient to set the rate of change in the logarithm of discrimination. The average value of – log *f* (*x*) is a common definition of information or entropy, and is related to many interpretations in terms of information theory (Cover & Thomas, 1991; Sims, 2018).

## Rotational invariance and Gaussian patterns

The scale, *x*, measures the perceptual difference between two entities or events. In some cases, the total difference, *x*, depends on the perceived differences along several distinct underlying dimensions. With two underlying dimensions, we may write

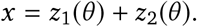

For a particular value of *x*, the parameter *θ* describes all of the combinations of the two underlying dimensions that add invariantly to *x*. If we let *x* = *r*^2^ and let the dependence of *z* on *θ* be implicit, we can write the prior expression equivalently as

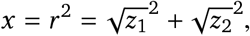

which defines a circle with coordinates along the positive and negative values of 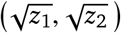, with a constant radius *r* that is rotationally invariant with respect to the parametric angle, *θ*. Traditionally, one uses 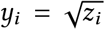, so that the radius, *r*, of a sphere has the familiar definition of a Euclidean distance

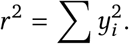

For each radial value, 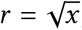, we can write {*y_i_* (*θ*)} as the sets indexed by the parameter *θ* for which the individual dimensional measures combine to the same invariant radius. If the angles of rotation with equivalent radius occur with equal probability or without prior bias, then radial values are rotationally invariant with respect to probability or prior likelihood.

I now show that rotational invariance leads to the Gaussian pattern as a special case of the general exponential form. In the exponential form derived in earlier sections, *λx* described the stretch-invariant perceptual scale. To express that scale in terms of a rotationally invariant radial measure, *r*, we note that *x* = *r*^2^ and we let *λ* = *πν*^2^. Thus, we can write the stretch-invariant incremental perceptual measure as

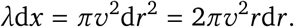

The general exponential form is

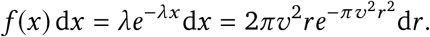

At a given radius, *νr*, if, by rotational invariance, all combinations of values for the underlying measurement dimensions occur without bias or prior information, then the total probability in a radial increment, *νdr*, is spread uniformly over the circumferential path with length 2*πνr*.

A radial vector intersects a fraction of the total probability density in the circumferential path in proportion to 1/2*πνr*. Thus, the probability along an increment *νdr* of the radial vector is

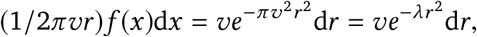

invariantly with respect to the angle of orientation of the radial vector. This expression is the Gaussian distribution, with *r*^2^ as the squared deviation from the mean or central location, and with parameters commonly written as *λ* = 1/2*σ*^2^ and 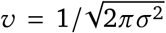 for variance *σ*^2^. The variance is simply the average value of the squared radial deviations, *r*^2^ = *x*.

We can also write the Gaussian in terms of the standard perceptual scale, *x*, as

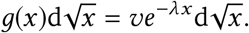

When we consider the standard perceptual scale, *x*, with respect to the incremental square-root scale, 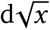, we obtain a Gaussian. The incremental square-root scale makes sense when we consider *x* as an aggregate measure of the sum of underlying perceptual dimensions. Each dimension naturally takes on a square-root scaling relative to the invariant total distance, because of the Euclidean measure of squared distance as the sum of squares along each dimension.

## Discussion

Any reasonable perceptual scale must satisfy the simple affine invariances of shift and stretch. I have shown that those invariances are sufficient to explain the exponential form of the universal law of generalization. I have also shown that an additional common invariance of rotation explains why some observed patterns of generalization follow a Gaussian rather than exponential pattern. The Gaussian pattern is, in fact, a special case of exponential scaling, when the scale is a squared Euclidean distance over several underlying dimensions.

Previous explanations also generate the exponential pattern of the universal law (Shepard, 1987; Chater & Vitányi, 2003; Sims, 2018). The reason those explanations succeed is that they include assumptions about shift invariance, which by itself generates an exponential pattern. All of the other assumptions and language associated with those prior explanations are superfluous with respect to the exponential form. Conclusions about rate of change in discrimination typically associate with an assumption about stretch invariance or, equivalently, conservation of average value.

It is certainly true that additional assumptions will lead to more precise predictions, which may then be tested to rule out particular mechanisms. But those additional assumptions and tests do not directly bear on the general exponential form itself.

I do not know of explicit prior explanations that unify the Gaussian pattern with the universal exponential law. Such explanations, if they exist, will generally reduce to the assumption of rotational invariance. Again, additional assumptions or arguments about particular underlying mechanisms are superfluous with regard to the general pattern.

It is, of course, interesting to consider what underlying perceptual mechanisms lead to the universal law. However, almost certainly, there is no single mechanism that could explain such a widely observed pattern. General patterns require general explanations that apply broadly. The simple invariances of meaningful measurement scales provide that general explanation for the observed patterns of perceptual scaling.

## Acknowledgments

National Science Foundation grant DEB–1251035 and the Donald Bren Foundation support my research. I completed this work while on sabbatical in the Theoretical Biology group of the Institute for Integrative Biology at ETH Zürich.

